# Skin Lesion Segmentation with Improved C-UNet Networks

**DOI:** 10.1101/382549

**Authors:** Hongda Jiang, Ruichen Rong, Junyan Wu, Xiaoxiao Li, Xu Dong, Eric Z Chen

## Abstract

—This paper proposes an innovative method for Part1, skin lesion segmentation of the ISIC 2018 Challenge. Our network C-UNet is based on UNet network, we combined several methods on this basic network which made some improvements on Jaccard Index ultimately, our method yield an average Jaccard Index of 0.77 on the On-line validation dataset.

## I. INTRODUCTION

Malignant melanoma is a common and threading skin cancer, and increasing year by year. It’s one of the most rapidly increasing cancers all over the world [1]. If the skin cancer is recognized earlier and treated surgically, the cure rate is much higher. Automatically segmenting melanoma from the surrounding skin is an essential step in computerized analysis of dermoscopic images [2] [3]. Dermatologists diagnose melanoma by visual inspections of mole using clinical assessment tools such as ABCD. However, computer vision tools have been introduced to assist in quantitative analysis of skin lesions. Due to the development of deep learning in the medical field, our team proposed an improved UNet segmentation network. A series of experiments done on the proposed method show that the proposed method delivers better accuracy and robust segmentation results.

## II. MATERIALS

### A. Database

For training, we employed the ISIC 2018 Challenge official dataset, with 2594 dermoscopic images and the corresponding lesion masks, we took two-third of the sample images as training sets, leaving one-third of the sample images as test sets. The lesion types involved include nevus, seborrhoeic keratosis and melanoma. The goal is to produce accurate binary masks of various skin lesions against a variety of background. Besides training set, the organizers provide a validation dataset that includes 100 images. The participants can submit the binary masks of these 100 images and evaluate the segmentation performance online. Additional test dataset with 1000 images is provided for final evaluation. The final rank is based on Jaccard Index.

### B. Data Augmentation

As the images with various dimensions. Images were first resized we tried 450×600 and 256×342,128× 172. In terms of data augmentation, founded through experiments that the network tends to be biased towards darker areas during training. The contrast between the lesion area and the non-lesion area is not very obvious. Therefore, we use the histogram equalization method to enhance color contrast of the image,so we combined this two images, and train the enhanced image with six channels in order to learn more features.

**Fig. 1.**
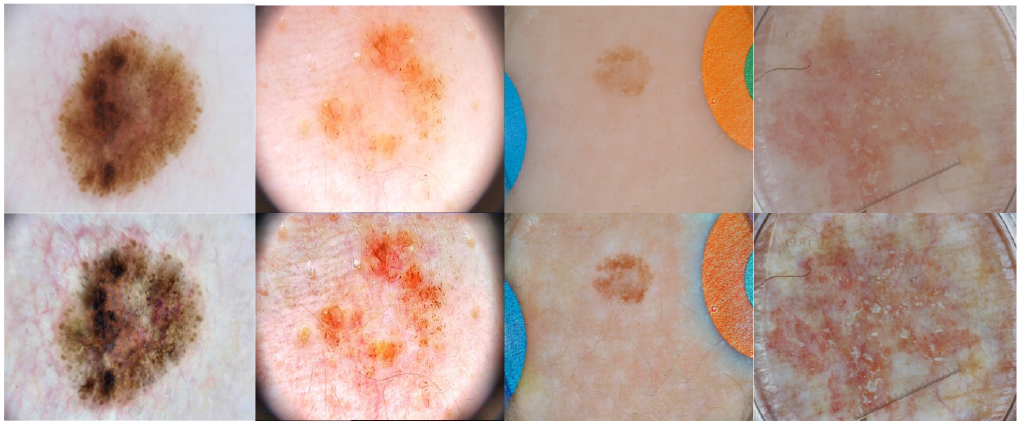
Example of lesion data augmentation. The first row are some original images, and the second row are some data-augmented images.

### C. Implementation

Our method was implemented with Python based on Tensorflow. The experiments were trained on two GPUs of Nvidia GeForce GTX 1080ti with 11GB memory per each and Intel (R) i7-7700k 4.2 GHz CPU.

## III. C-UNet NETWORKS

Our segmentation neural network is designed based on UNet [4]. According to the training results, we made some improvement on the UNet and finally C-UNet network is obtained. In the beginning, we used the UNet network structure for training. The jaccard index result is around 0.72 (If jaccard index<0.65, let jaccard index=0), and the experimental results showed that some generated mask was not fully connected, but the ground truth of each example is completely and connected, so we consider changing the size of the receptive field. A new version of UNet2 use a multi-scale convolution like inception v3 [5] in the encoding part, as it shown in TABLE I. And we use dilated convolution block before the pooling layer, the same thing between the dilated convolution [6] and the ordinary convolution is that the size of the convolution kernel is the same, and have the same number of parameters but the dilated convolution has a larger receptive field. In our C-UNet we choose the dilated coefficient as 4. The experimental results showed a significant improvement, jaccard index reached about 0.74, and still let jaccard index=0 when jaccard index <0.65. Secondly, in order to better utilize the features learned by different layers, we adopt the structure of RNN [7] in front of each pooling layer, and realize the fusion of features in different layers through the function of circular convolution which informed UNet3. The final jaccard index result is around 0.752. Finally, we found that the final result of the validation dataset, there are some jaccard<0.65 prediction results, these bad prediction inference the final average results, so we refer to the Deeplab network [8], adding image segmentation post processing, the mask generated by the network is refined using the conditional random field (CRF), at last the final validation result is maintained at 0.77, and the test result is maintained at 0.755. The result was shown in TABLE IV.

**TABLE 1.** MULTISCALE CONVOLUTION BLOCK

|  | Layer | Filter Size | Stride | Dilation |
| --- | --- | --- | --- | --- |
| Conv_Block | Multi_Conv | 5×5 | 1 |  |
|  |  | 3×3 | 1 |  |
|  |  | 1×1 | 1 |  |
|  |  | 1×7,7×1 | 1 |  |
|  | RCL | 3×3 | 1 |  |
|  |  | 3×3 | 1 |  |
|  |  | 3×3 | 1 |  |
|  | Dilated_Conv | 3×3 |  | 4 |

**TABLE 2.** DECONVOLUTION BLOCK

|  | Layer | Filter Size | Stride |
| --- | --- | --- | --- |
| Deconv_Block | Deconvolution | 3×3 | 2 |
|  | Concatenet |  |  |
|  | Conv | 3×3 | 1 |
|  | Conv | 3×3 | 1 |
|  | RCL | 3×3 | 1 |
|  |  | 3×3 | 1 |
|  |  | 3×3 | 1 |

**TABLE 3.** C-UNet Architecture

| Layer | Filter size | Stride | NO. of features |
| --- | --- | --- | --- |
| Conv_Block1 |  |  | 32 |
| Pooling | 3×3 | 1 | 32 |
| Conv_Block2 |  |  | 64 |
| Pooling | 3×3 | 1 | 64 |
| Conv_Block3 |  |  | 128 |
| Pooling | 3×3 | 1 | 128 |
| Conv_Block4 |  |  | 256 |
| Pooling | 3×3 | 1 | 256 |
| Conv_Block5 |  |  | 512 |
| Pooling | 3×3 | 1 | 512 |
| Conv_Block6 |  |  | 1024 |
| Conv_Block7 |  |  | 1024 |
| Deconv_Block1 |  |  | 512 |
| Deconv_Block2 |  |  | 256 |
| Deconv_Block3 |  |  | 128 |
| Deconv_Block4 |  |  | 64 |
| Deconv_Block5 |  |  | 32 |
| Conv | 1×1 | 1 | 1 |
| Crf |  |  | 1 |

**TABLE 4.** JACCARD RESULT OF DIFFERENT ARCHITECTURE

| Architecture | Jaccard Index |
| --- | --- |
| UNet | 0.72 |
| UNet2 | 0.74 |
| UNet3 | 0.752 |
| C-UNet | 0.755 |

### A. Training

We train the network using Adam optimization [9] in the beginning with batch size of 8, and use SGD optimization [10] for finetune. The initial learning rate is set as 0.0001, exponential decay method has been used on learning rate after one epoch. In order to reduce overfitting, we use dropout with p = 0.9 in each convolution layer. As for loss function, we choose the cross-entropy loss as the training loss function in the beginning, and choose the dice loss for finetune, we choose different weights on the foreground dice loss and background dice loss as it shown in Equations 3. As shown in Equations below, where y^i^ is the label for each sample, ŷ^i^ is the predicted label, and X and Y in Equation 2 represent the predicted mask and ground truth, respectively.

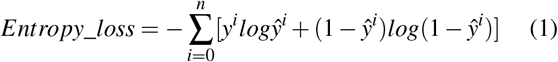

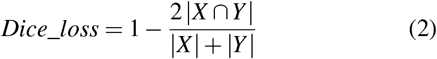

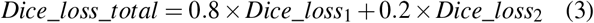

### B. Result

To show how our proposed method works on different images, we show our segmentation results of some challenging images in Fig. 2 . Images with training image, probability map, ground truth and segmentation result are shown in the first column, second column, third column and last column of Fig. 2, respectively. From Fig. 2, it can be seen that our method works very well in most of these challenging images. For the test dataset the overall average performance are derived as 91.8% Accuracy. The overall Jaccard Index score is 0.755, Dice coefficient score is 0.82.

## IV. Related Work

Some other techniques are implemented but lead to lower performance.

We replaces the original UNet encoder with rich-feature models like ResNet-101 [11], VGG16 [12] and use pretrained weights from ImageNet and only fine-tune the decoder parts. Compared to vanilla UNet encoder, ResNet and VGG are supposed to abstract complicated features under different resolution. So we expect using these backbones as encoder could lead to higher performance on complicated images. But to our surprise, the Jaccard Index drops dramatically. Also adding border weights to the UNet loss function results in slightly lower scores.

Detection based algorithm like Mask-RCNN [13] are also tested on this task. A Mask-RCNN model with pretrained ResNet-101 backbone on COCO dataset gives 0.65 Jaccard Index on the validation dataset. One of the biggest challenge with Mask-RCNN is to combine small pieces of predicted regions into one big mask. We reduce the number of predicted regions and allow very low threshold in inference in order to keep very large bounding box. Otherwise, only small regions with high confidence scores are kept by the model. Then NMS is used to merge all bounding-boxes into one large bounding box. The performance is similar to vanilla UNet in majority of the easy images but still meet difficulties to combine predicted regions in challenging images.

Some post-processing tricks are also implemented. We first use Dilation + Erosion to smooth the border of masks. Then we remove small holes and small objects outside of the main masks. While the above two steps are not necessary for majority of images and have no influence on the scores. Watershed algorithm is also tried to enlarge mask region in certain images. But adding this post-processing steps will lead to unpredictable results for other images, so it’s not included in our final model.

**Fig. 2.**
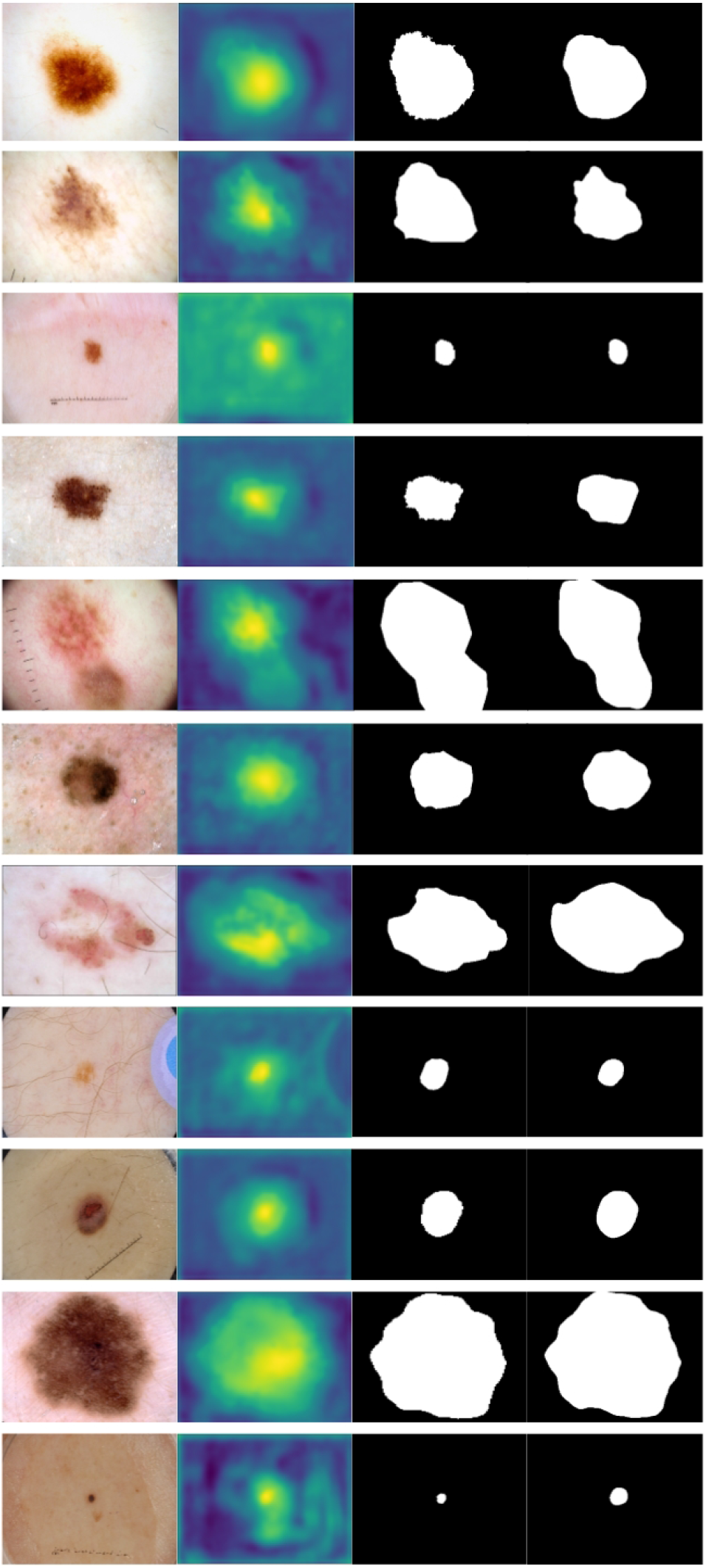
Datasets and qualitative results of our experiments. From left to right: image, probability map, ground truth, segmentation result.

